## Supplemental files-Extended data for "Phenotypic plasticity in bacterial elongation among closely related species"

### Extended Data Figure 1

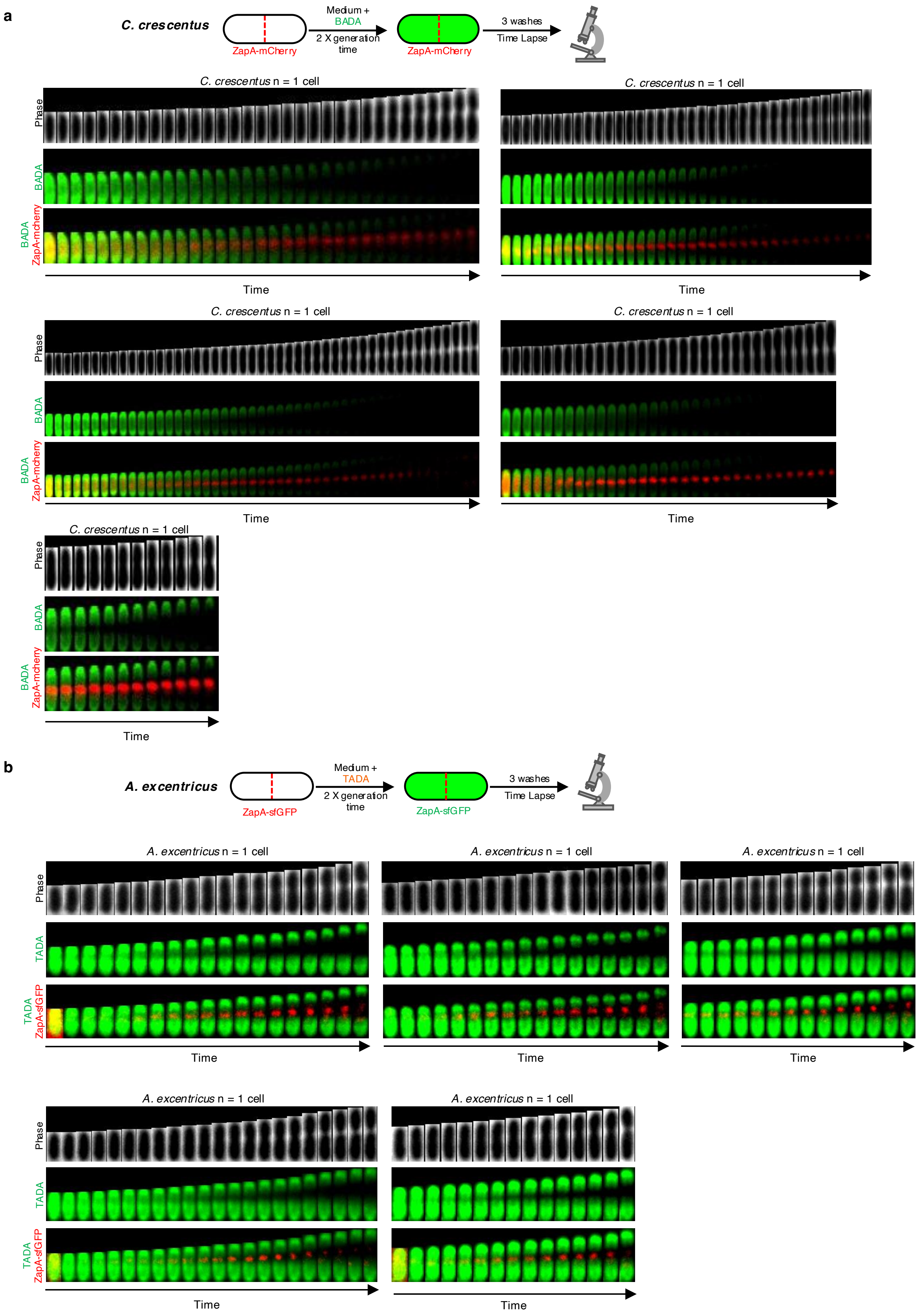

**Extended Data Fig. 1. *C. crescentus* and *A. excentricus* have different patterns of PG synthesis**

(**a**) Schematic of the pulse-chase experiment using the FDAA BADA (green) in *C. crescentus* cells expressing ZapA-mCherry (red). Kymographs from five different cells from this pulse-chase experiment are shown.

(**b**) Schematic of the pulse-chase experiment using the FDAA TADA (green) in *A. excentricus* cells expressing ZapA-sfGFP (green). Kymographs from five different cells from this pulse-chase experiment are shown.

#### Extended Data Figure 2

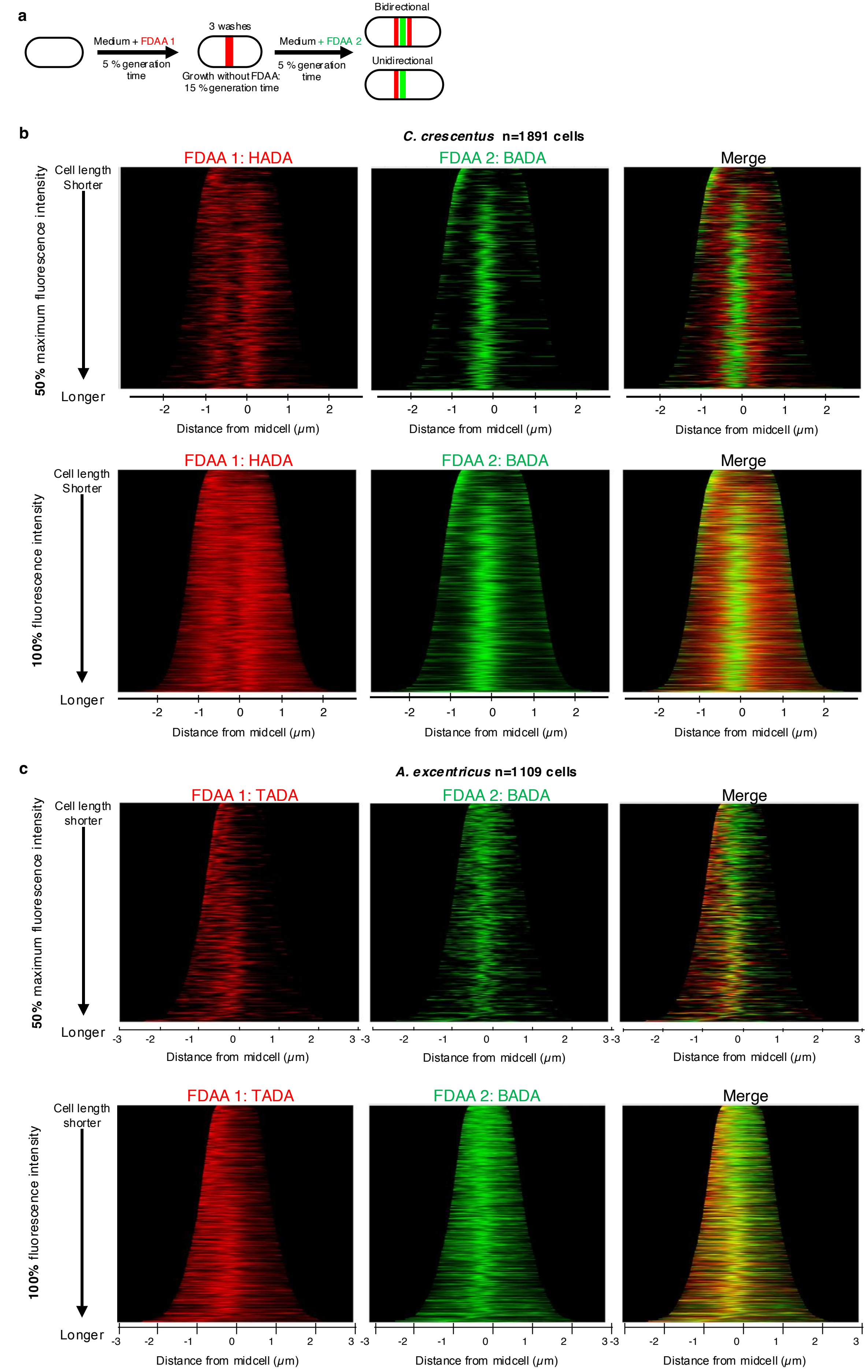

**Extended Data Fig 2.** Demographs from dual short-pulse labeling with FDAAs in *C. crescentus* and *A. excentricus*.

**(a)** Schematic depicting the dual short-pulse experiment. Cells were first labeled with one FDAA for 5% of their generation time (HADA for *C. crescentus* and TADA for *A. excentricus*), washed with PYE to remove free FDAA, allowed to grow for 15% of their generation time and then labeled with a second FDAA (BADA) for 5% of their generation time, washed again, and imaged with phase and fluorescence microscopy.

(**b-c**) Demographs showing the fluorescence intensities of both FDAA signals in (**b**) *C. crescentus* cells and (**c**) *A. excentricus* cells. Demographs are presented for each FDAA independently, as well as together on the same graph. Cells were arranged by length with the maximum fluorescence intensity of the FDAA signal to the left. 50% of the maximum fluorescence intensities (*top*) and 100% of the fluorescence intensities (*bottom*) are shown. The first FDAA is represented in red and the second FDAA in green.

### Extended Data Figure 3

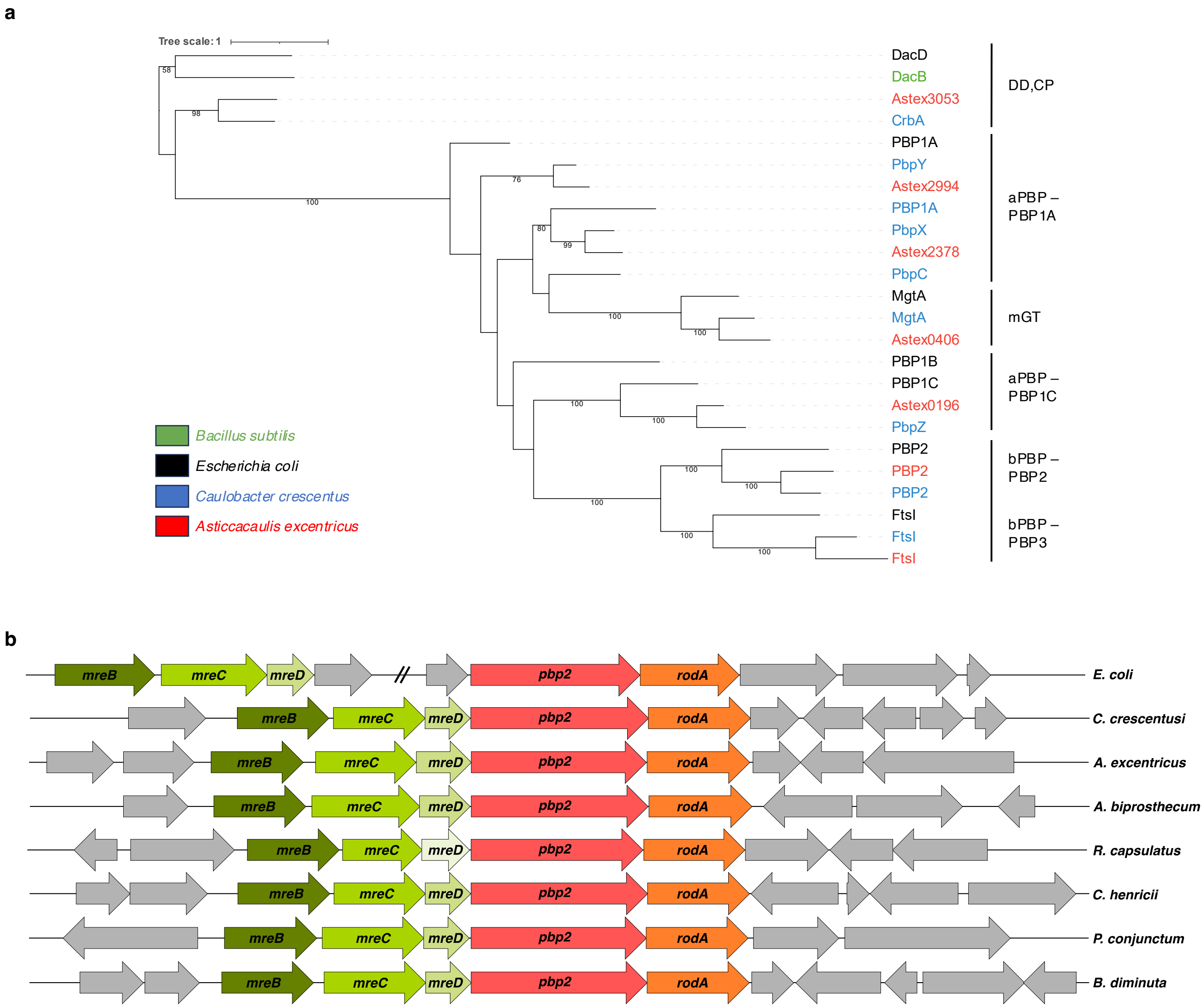

**Extended Data Fig. 3. Sequence alignment of PBPs from different species and genomic organization of the *pbp2* and *mreB* genes.**

**(a)** Phylogenetic tree of PBP sequences from *E. coli*, *B. subtilis*, *C. crescentus* and *A. excentricus*. Each species is represented by a specific color: black for *E. coli*, blue for *C. crescentus*, red for *A. excentricus*, and green for *B. subtilis*. aPBP: class A penicillin-binding protein; bPBP: class B penicillin-binding protein; D,D-CP: D,D-carboxypeptidase; and mGT: mono-glycosyltransferase. Bootstrap values >50 % are indicated at their respective nodes (based on 100 replicates). The final tree was formatted using iTol[73].

**(b)** Overview of the genomic organization of *pbp2* and *mreB* genes in selected species. The figure displays the genomic arrangement of the *pbp2* and *mreB* loci across the species of interest, presented from top to bottom: *E. coli*, *C. crescentus*, *A. excentricus*, *A. biprosthecum*, *R. capsulatus*, *C. henricii*, *P. conjunctum*, and *B. diminuta*. The genes are color-coded according to their type: *pbp2* is represented in red, and *rodA* in orange. *mreB*, *mreC*, and *mreD* genes are shown in shades of green, with *mreD* from *R. capsulatus* depicted in a lighter green to indicate that it did not meet the identity and e-value cut-off. All other neighboring genes are shown in gray to indicate non-relevant or less conserved regions. The length of each arrow and the spacing between them reflect the gene sizes and intergenic distances in nucleotides.

### Extended Data Figure 4

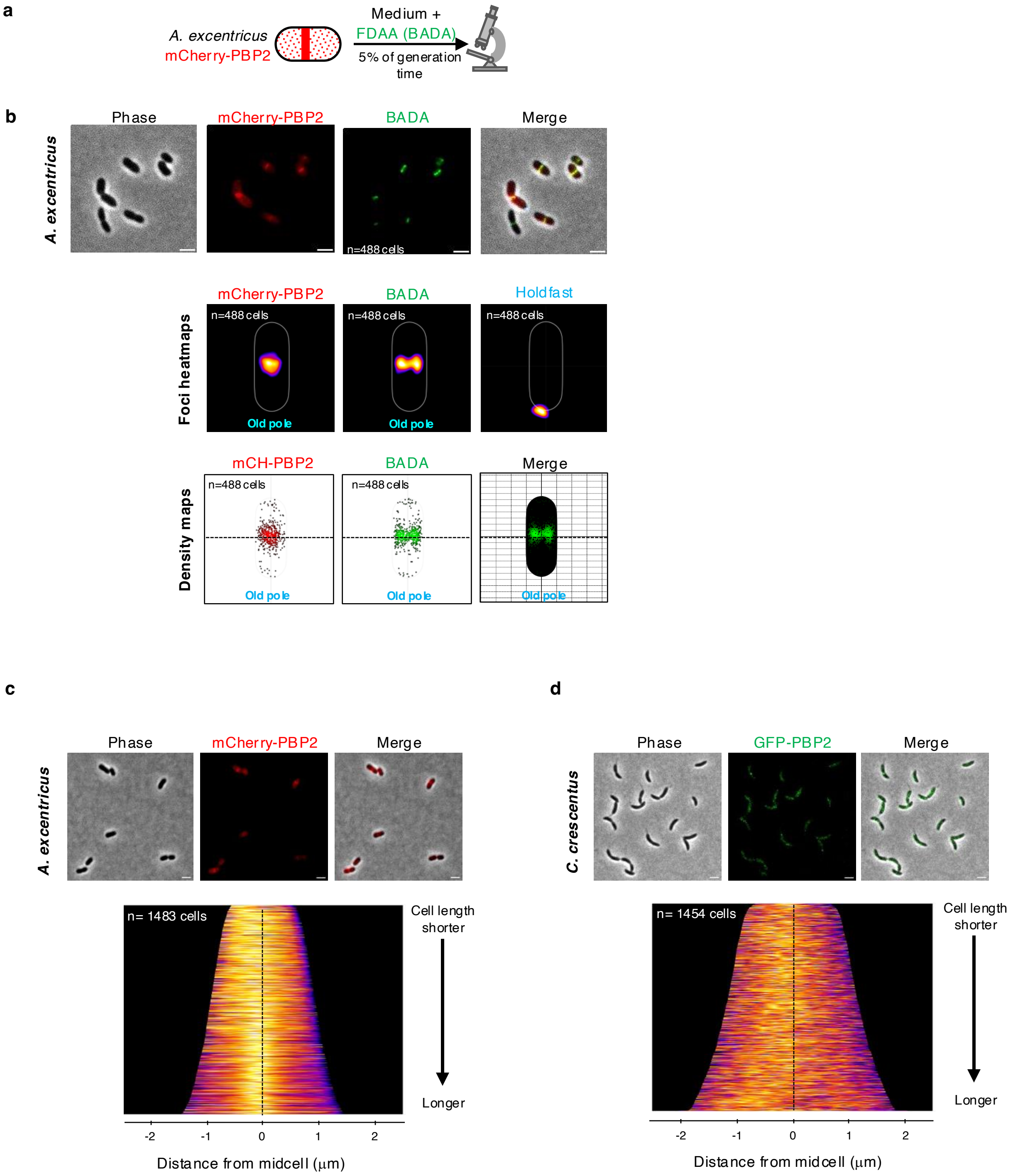

**Extended Data Fig. 4. PBP2 localizes close to the midcell in *A. excentricus* while PBP2 is dispersed in *C. crescentus*.**

**(a)** Short-pulse FDAA (BADA) labeling of *A. excentricus* cells expressing mCherry-PBP2. A schematic depicting the short-pulse experiment is shown. Cells were labeled with 500 µM BADA for 5% of their generation time, fixed with 70% ethanol and imaged.

**(b)** *Top:* Representative images of the short-pulse labeled *A. excentricus* mCherry-PBP2 cells are shown. Left to right: Phase channel, mCherry-PBP2, BADA, and merged images with WGA labeling. Scale bar: 2 µm. *Middle:* Heatmaps of mCherry-PBP2 and BADA foci at the population level are shown. *Bottom:* Density maps of mCherry-PBP2 and BADA foci at the population level are shown, with the black line indicating the midcell.

**(c)** Subcellular localization of mCherry-PBP2 in *A. excentricus*. *Top:* Representative images are shown with phase, mCherry-PBP2, and merged images. Scale bar: 2 µm. *Bottom:* A demograph showing the localization of the fluorescence intensity of mCherry-PBP2 at the population level, with each cell oriented such that the pole with the maximum fluorescence intensity is to the left.

**(d)** Subcellular localization of GFP-PBP2 in *C. crescentus*. *Top:* Representative images are shown with phase, GFP-PBP2, and merged images. Scale bar: 2 µm. *Bottom:* A demograph showing the localization of the fluorescence intensity of GFP-PBP2 at the population level, with each cell oriented such that the pole with the maximum fluorescence intensity is to the left.

### Extended Data Figure 5

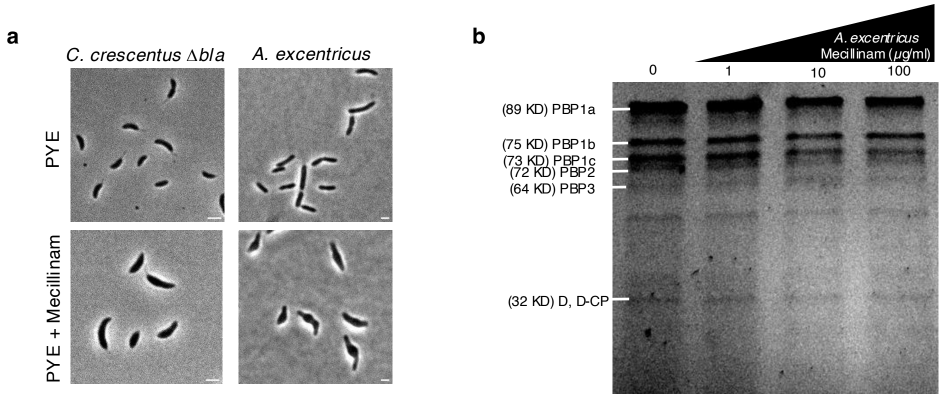

**Extended Data Fig. 5. Mecillinam targets PBP2 in *C. crescentus* and *A. excentricus***

(**a**) Phase contrast images of the Δ*bla* β-lactam-sensitive strain of *C. crescentus*, and *A. excentricus* WT cells treated with or without mecillinam (50 μg $ml^{-1}$).

(**b**) SDS-PAGE gel image for mecillinam titration against the different PBPs in *A. excentricus.* Whole cells were treated with various concentrations of mecillinam and subsequently labeled with Boc-FL prior to SDS-PAGE.

### Extended Data Figure 6

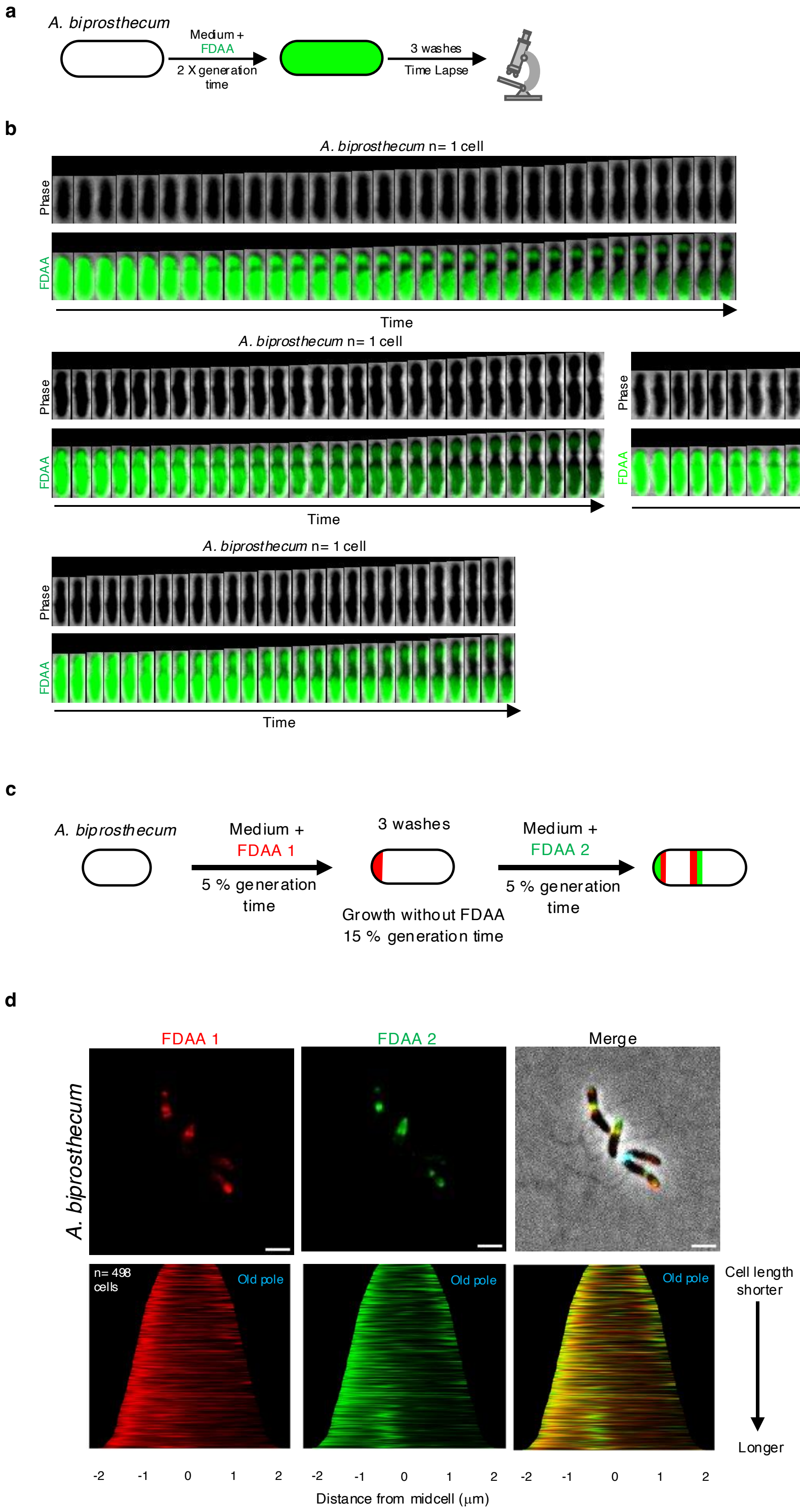

**Extended Data Fig. 6. Pulse-chase and dual short-pulse FDAA labeling experiments reveal that *A. biprosthecum* shows both polar and unidirectional midcell growth.**

(**a**) Schematic of the pulse-chase experiment. Whole-cell PG was labeled with 500 µm FDAA (TADA, green) over two generations, followed by washes with PYE to remove free FDAA from the medium. Subsequent growth in the absence of FDAA was followed by time-lapse microscopy. During this chase period, the loss of FDAA signal corresponds to new PG synthesis/turnover.

(**b**) Representative cells from the pulse-chase experiment. Five different *A. biprosthecum* cells and corresponding kymographs are shown.

**(c)** Schematic depicting the dual short-pulse experiment. Cells were first labeled with one FDAA (TADA, red) for 5% of their generation time, washed with PYE to remove free FDAA, allowed to grow for 15 % of their generation time, and then labeled with a second FDAA (BADA, green) for 5% of their generation time, washed again, and imaged with phase and fluorescence microscopy.

**(d)** *Top:* Representative images showing each FDAA individually or merged with phase and old-pole labeling with WGA (cyan). Scale bar: 2 µm. *Bottom:* Demographs of the FDAAs showing the full range of their fluorescence signals, individually or merged in the same graph. Cells were arranged by length with the old pole (labelled with WGA, not shown) to the right.

### Extended Data Figure 7

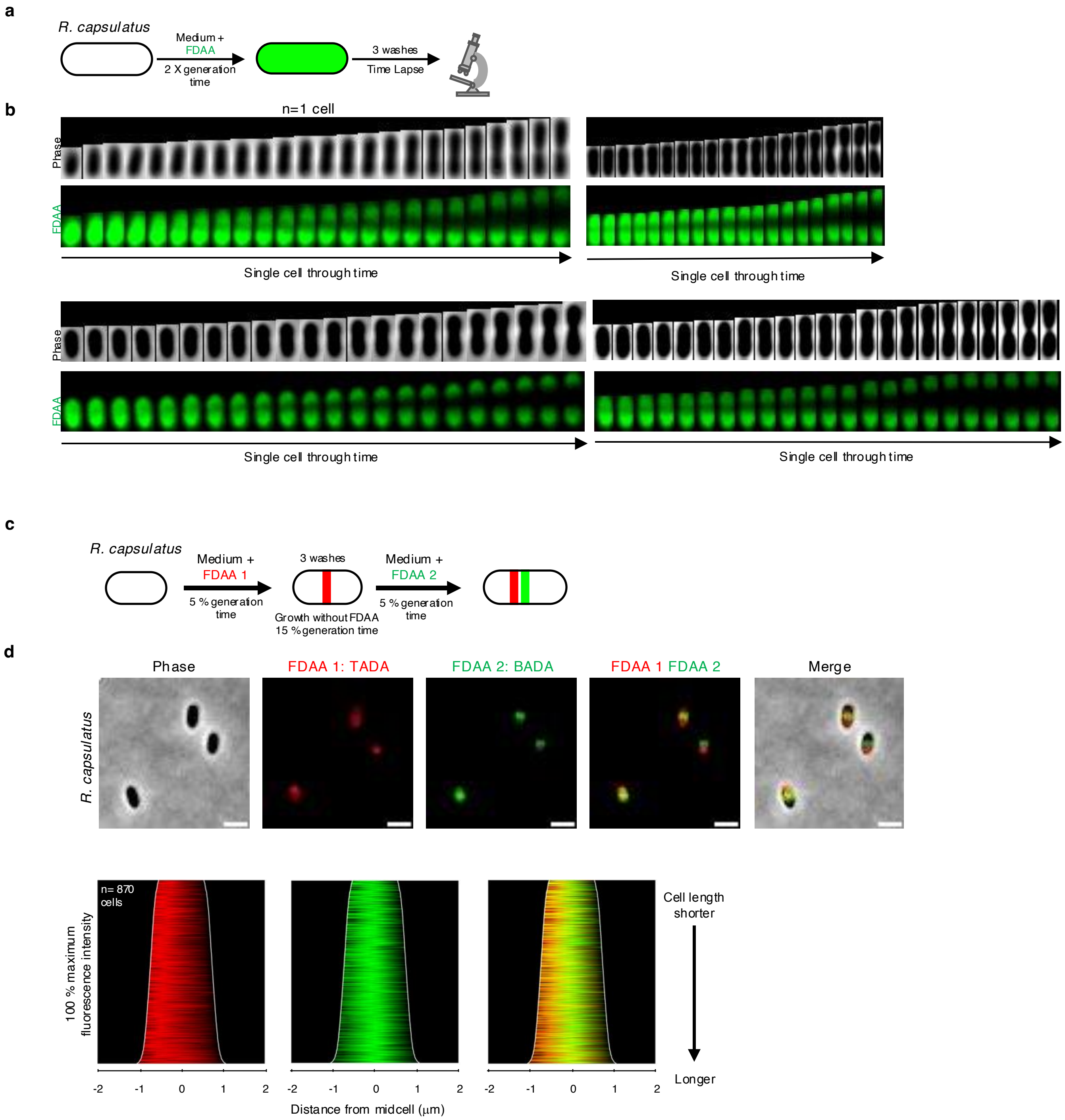

**Extended Data Fig. 7. Pulse-chase and dual short-pulse FDAA labeling experiments reveal that *R. capsulatus* shows unidirectional midcell growth.**

(**a**) Schematic of the long-pulse and chase experiment. Whole-cell PG was labeled with 250 µm FDAA (TADA, green) over two generations, followed by washes with PYS to remove free FDAA from the medium. Subsequent growth in the absence of FDAA was followed by time-lapse microscopy. During the chase period, the loss of FDAA signal corresponds to new PG synthesis/ turnover.

(**b**) Representative cells during from the pulse-chase experiment. Four different *R. capsulatus* cells and corresponding kymographs are shown.

**(c)** Schematic depicting the dual short-pulse experiment. Cells were first labeled with one FDAA (TADA, red) for 5% of their generation time, washed with PYS to remove free FDAA, allowed to grow for 15% of their generation time and then labeled with a second colored FDAA (BADA, green) for 5% of their generation time, washed again, and imaged with phase and fluorescence microscopy.

**(d)** *Top:* Representative images showing each FDAA individually or merged with phase and old-pole labeling with WGA (cyan). Scale bar: 2 µm. *Bottom:* Demographs of the FDAAs showing the full range of their fluorescence signals, individually or merged in the same graph. Cells were arranged by length with the maximum fluorescence intensity to the left.

### Extended Data Figure 8

**a**

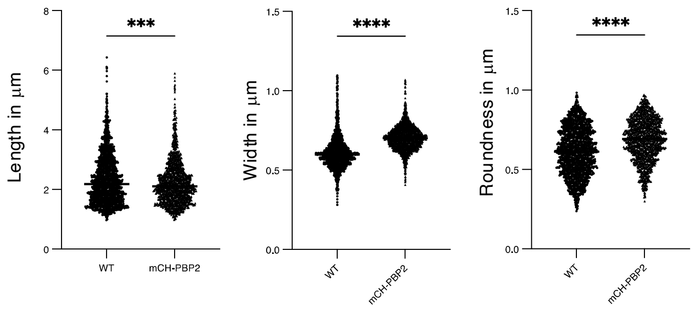


**b**

| Strain | Mutation |
|---|---|
| mCherry-PBP2 | FTSW : L34V and L35V |

**c**

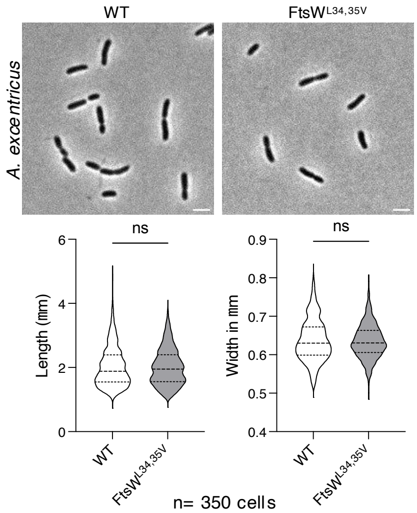


**d**

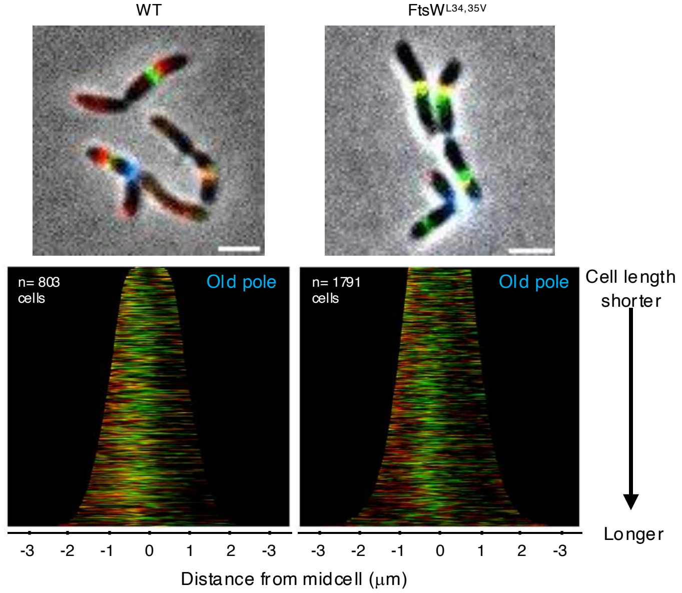

**Extended Data Fig. 8. Validation of the mCherry PBP2 fusion in *A. excentricus*.**

(**a**) Graphs comparing the length, width, and roundness of cells expressing the mCherry-PBP2 fusion, compared to WT. Statistical analysis was performed using ANOVA with Dunnett's correction for multiple comparisons, with significance indicated as ***P < 0.001 and ****P < 0.0001, relative to WT.

(**b**) Results of whole-genome sequencing for strain *mCherry-PBP2*. The *mCherry-pbp2* strain carries two missense mutations (L34V and L35V) in the *ftsW* gene.

(**c**) Phenotype of the *ftsW* L34V and L35V mutations backcrossed into WT. Representative phase-contrast images and graphs comparing the length and width of *ftsW* $^{L34V\ L35V}$to WT (n=350 cells). Scale: 2 μm.

(d) *Top:* Representative images are shown. *Left:* merge image of FDAA 1 TADA, FDAA 2 BADA, and phase contrast of WT cells. *Right:* merge image of FDAA 1 TADA, FDAA 2 BADA, and phase contrast of *ftsW* $^{L34V\ L35}$ cells. Scale bars: 2 μm. *Bottom:* Demographs showing the fluorescence intensity of both FDAA signals in WT and *ftsW* $^{L34V\ L35V}$ cells. Cells were arranged by length with the old pole (labelled with WGA, not shown) to the right. 50% of the maximum fluorescence intensities are shown.

**Extended Data Text 1:**

To assess the functionality of our mCherry-PBP2 fusion in *A. excentricus*, we compared the cell dimensions of the strain to WT, in terms of length, width, and roundness. Cells expressing the mCherry-PBP2 fusion showed a slight increase in width and roundness compared to WT cells, as expected for a slight reduction in PBP2 activity[74] (**Extended Data Fig. 8a**). We sequenced the genomic DNA of the strain and found that the *mcherry-pbp2* strain harbored two missense mutations in the *ftsW* gene encoding the glycosyltransferase implicated in division (**Extended Data Fig. 8b)**. To ensure that these mutations in *ftsW* did not affect any observed phenotypes, we conducted backcross experiments to transfer these mutations into the WT background. The resulting strain showed no significant differences in phenotype compared to WT strains in phase contrast microscopy or FDAA dual short-pulse analyses (**Extended Data Fig. 8c-d)**.

### Tables

#### Table S1: Strains used in the study

| Strain | Description and or genotype | Reference or source |
|---|---|---|
| ***E. coli*** | | |
| NEB 5-alpha | *fhuA2Δ(argF-lacZ)U169 phoA glnV44 Φ80Δ(lacZ)M15 gyrA96 recA1 relA1 endA1 thi-1 hsdR17* | NEB |
| YB 8804 | DH5α / pGFPC-5-$zapA_{ex}$ | This study |
| YB 8873 | *E. coli* Top10/ pCHYC-4-$zapA_{cc}$ | [29] |
| YB 9852 | DH5α / pNPTS139-*mch-Aex-pbp2* | This study |
| YB 9856 | DH5α / pNPTS138-*Aex-ftsW** | This study |
| ***C. crescentus*** | | |
| YB 135 | Wild-type strain CB15 | [53] |
| YB 127 | Wild-type strain CB15N | [75] |
| YB 8872 | CB15N *zapA::zapA-mcherry* | [76] |
| YB 9460 | CB15N *pbp2::gfp-pbp2* | [33] |
| CS 606 | CB15N Δ*bla* | [77] |
| ***A. excentricus*** | | |
| YB 258 | *A. excentricus* AC48 | [78] |
| YB 8871 | AC48 *zapA::zapA-gfp* | This study |
| YB 8806 | AC48 *pbp2::mcherry-pbp2* | This study |
| YB 8807 | AC48 *zapA::zapA-gfp pbp2::mcherry-pbp2* | This study |
| YB 9858 | AC48 *ftsW** | This study |
| ***Other species*** | | |
| YB 3934 | *Caulobacter henricii* | Brun Lab UdeM |
| YB 5193 | *Brevundimonas diminuta* | Brun Lab UdeM |
| YB 7710 | *Phenylobacterium conjunctum* | Brun Lab UdeM |
| YB 642 | *Asticcacaulis biprosthecum* | [79] |
| YB 4650 | *Rhodobacter capsulatus SB1003* | Bauer Lab[54] |

**Table S2: Plasmids used in the study**

| Plasmids | | Antibiotic | References |
|---|---|---|---|
| pNPTS139 | sacB-containing Litmus 39 derivative suicide vector used for double homologous recombination | Kan | M.R.K. Alley |
| pNPTS138 | sacB-containing Litmus 38 derivative suicide vector used for double homologous recombination | Kan | M.R.K. Alley |
| pGFPC-5 | Integration plasmid used for creating C-terminal fusions to eGFP | Spec/Strep | [80] |
| pCHYC-5 | Integration plasmid used for creating C-terminal fusions to mCherry | Spec/Strep | [80] |
| pCHYC-4 | Integration plasmid used for creating C-terminal fusions to mCherry | Gent | [80] |
| psfGFPC-5-$zapA_{ex}$ | pGFPC-5 bearing the C-terminal fragment of *zapA* (*astex_1642*), with the *egfp* gene replaced by *sfgfp* | Spec/Strep | This study |
| pCHYC-5-$zapA_{ex}$ | pCHYC-5 bearing the C-terminal fragment of *zapA* (*astex_1642*) | Spec/Strep | This study |
| pCHYC-4-$zapA_{cc}$ | pCHYC-4 bearing the C-terminal fragment *zapA* (*ccna_03356*); | Gent | [76] |
| pNTPS139-*mCherry-PBP2* | pNTPS derivative for in-frame replacement of *pbp2* (*astex_1631*) with *mCherry-pbp2* | Kan | This study |
| pNTPS138-*Aex-ftsW** | pNTPS derivative for backcrossing *ftsW* mutation | Kan | This study |

**Table S3: Genome IDs and mode of cell elongation**

| Strain | Taxonomy ID | Mode of cell elongation |
|---|---|---|
| *Agrobacterium tumefaciens* | 176299 | Polar[9] |
| *Asticcacaulis aquaticus* | 2984212 | Binary fission[81] |
| *Asticcacaulis benevestitus* | 1121022 | Binary fission[82] |
| *Asticcacaulis biprosthecum* | 76891 | Polar plus unidirectional midcell elongation – This work |
| *Asticcacaulis excentricus* | 573065 | Unidirectional midcell elongation - this work |
| *Brevundimonas diminuta* | 751586 | Bidirectional midcell elongation - this work |
| *Brevundimonas naejangsanensis* | 588932 | Unknown |
| *Brucella abortus* | 235 | Polar[9] |
| *Caulobacter crescentus* | 565050 | Bidirectional midcell elongation – this work |
| *Caulobacter fusiformis* | 69396 | Binary fission[83] |
| *Caulobacter henricii* | 69395 | Bidirectional midcell elongation – this work |
| *Caulobacter segnis* | 509190 | Unknown |
| *Escherichia coli* | 511145 | Dispersed[27] |
| *Henriciella marina* | 1121949 | Binary fission[84] |
| *Hirschia baltica* | 582402 | Budding[85] |
| *Hyphomicrobium denitrificans* | 582899 | Budding[86] |
| *Hyphomonas neptunium* | 228405 | Budding[87-89] |
| *Maricaulis maris* | 394221 | Binary fission[90] |
| *Oceanicaulis alexandrii* | 1122613 | Binary fission[91] |
| *Peiella sedimenti* | 3061083 | Binary fission[92] |
| *Phenylobacterium composti* | 457173 | Unknown |
| *Phenylobacterium conjunctum* | 1298959 | Unidirectional midcell elongation - this work |
| *Phenylobacterium immobile* | 21 | Unknown |
| *Phenylobacterium koreense* | 266125 | Unknown |
| *Phenylobacterium zucineum* | 450851 | Unknown |
| *Rhodobacter capsulatus* | 1060 | Unidirectional midcell elongation - this work |
| *Rhodobacter sphaeroides* | 557760 | Bidirectional midcell elongation[49] |
| *Rhodomicrobium vannielii* | 648757 | Budding[87,88] |
| *Rhodopseudomonas palustris* | 1076 | Budding[87,88] |
| *Robiginitomaculum antarcticum* | 1123059 | Binary fission[93] |
| *Sagittula stellata* | 388399 | Budding ?[9]- No primary reference |
| *Sinorhizobium meliloti* | 382 | Polar[9] |
